# CD39 is expressed on functional effector and tissue resident memory CD8+ T cells

**DOI:** 10.1101/2024.03.15.585252

**Authors:** Jordan F. Isaacs, Hanna N. Degefu, Tiffany Chen, Sierra A. Kleist, Shawn C. Musial, Myles A. Ford, Tyler G. Searles, Chun-Chieh Lin, Alexander G. J. Skorput, Keisuke Shirai, Mary Jo Turk, George J. Zanazzi, Pamela C. Rosato

**Affiliations:** Department of Microbiology and Immunology, Geisel School of Medicine at Dartmouth College, Lebanon, NH, USA; Department of Pathology and Laboratory Medicine, Dartmouth Health, Lebanon NH, USA; Department of Neurology, Dartmouth Health, Lebanon NH, USA; Department of Medicine, Dartmouth Health, Lebanon NH, USA

## Abstract

The ecto-ATPase CD39 is expressed on exhausted CD8+ T cells in chronic viral infection and has been proposed as a marker of tumor-specific CD8+ T cells in cancer, but the role of CD39 in an effector and memory T cell response has not been clearly defined. We report that CD39 is expressed on antigen-specific CD8+ short-lived effector cells (SLECs), while it’s co-ecto-enzyme, CD73, is found on memory precursor effector cells (MPEC) *in vivo*. Inhibition of CD39 enzymatic activity during *in vitro* T cell priming enhances MPEC differentiation *in vivo* after transfer and infection. The enriched MPEC phenotype is associated with enhanced tissue resident memory (T_RM_) establishment in the brain and salivary gland following an acute intranasal viral infection, suggesting that CD39 ATPase activity plays a role in memory CD8+ T cell differentiation. We also show that CD39 is expressed on human and murine T_RM_ across several non-lymphoid tissues and melanoma, while CD73 is expressed on both circulating and resident memory subsets in mice. In contrast to exhausted CD39+ T cells in chronic infection, CD39+ T_RM_ are fully functional when stimulated *ex vivo* with cognate antigen. This work further expands the identity of CD39 beyond a T cell exhaustion marker.

## Introduction

CD8+ T cells play a crucial role in the immune response by killing infected cells and developing immunologic memory to rapidly clear pathogens upon re-infection^1^. During infection, anti-viral CD8+ T cells expand and differentiate broadly into memory precursor effector cells (MPEC) or short-lived effector cells (SLEC), which are defined by their expression of CD127/IL-7R and KLRG1^2,3,4^. Ultimately, MPECs give rise to subsets of memory CD8+ T cells; central memory (T_CM_), effector memory (T_EM_) and resident memory (T_RM_), which are defined by their circulation patterns^5,6^. T_RM_ have been identified in several non-lymphoid tissues in mice and humans and are characterized by their lack of recirculation and expression of CD69 and CD103^6–11^. T_RM_ are critical first line responders in tissues and can respond rapidly to pathogen re-encounter^6–11^.

CD8+ T cell memory differentiation is complex and, among other factors, controlled by several environmental cues, including sensing of extracellular metabolites like ATP and adenosine (ADO)^12–14^. Extracellular ATP (eATP) and ADO are potent immune modulators, controlling the activation or suppression of T cells through P2 (ATP) and P1 (ADO) receptors^15,16^. ATP can be released from damaged or dying cells, which acts as a danger signal and boosts the immune response via P2X and P2Y receptors^16,17,18^. To prevent excessive tissue damage, the immune system can fine-tune the delicate balance of eATP and ADO through the surface expression of purinergic ecto-enzymes CD39 and CD73^15^. CD39 hydrolyzes eATP to extracellular AMP (eAMP), which then requires CD73 to generate ADO^18^. Therefore, CD39 and CD73 are critical ‘immunological levers’ that can shift ATP-driven inflammation to ADO-mediated suppression^17^.

CD39 is expressed on exhausted CD8+ T cells responding to persistent antigen in chronic viral infection or tumors^19,20,21,22,23^. In individuals with HIV and HCV, CD39 expression on circulating virus-specific T cells correlated with increased viral load and reduced functionality^19^. In line with this association of CD39+ T cells and chronic antigen, CD39 has recently been identified as a marker of tumor-specific CD8+ T cells in several human tumors^20,22,24–27,28^. We and others have shown that tumor infiltrating lymphocytes (TILs) can have specificity for cancer unrelated antigens and thus CD39 may allow discrimination of tumor-specific T cells from these bystander T cells^20,29^.

Despite our understanding of CD39 in T cell exhaustion, its role in CD8+ T cell differentiation during an acute anti-viral response and throughout immunologic memory has not been fully characterized. CD8+ T cell differentiation is dictated in part by the strength of T cell receptor (TCR) stimulation, which can be modulated through eATP and ADO^30–32^. Therefore, CD39 and its co-ecto-enzyme CD73 have been proposed to play a role in T cell differentiation by enhancing ADO production and dampening TCR signaling strength^15^. CD39 and CD73 have been shown to have opposing expression patterns on activated T cells, where CD39 is increased and CD73 is decreased upon activation^18^. In addition, CD39 and CD73 are co-expressed on CD8+ T cells in the parenchyma of several non-lymphoid tissues.^33^ Despite these previous studies, the precise characterization of CD39 and CD73 co-expression on distinct T cell effector, circulating, and resident memory populations has not been established. To address these questions, we assessed CD39 and CD73 expression on antigen-specific CD8+ T cells in response to an acute viral infection. We found that CD39 was highly expressed on short-lived effector cells (SLEC) in contrast to memory precursor effector cells (MPEC), which expressed low CD39 and high CD73, suggesting ecto-enzyme expression may impact T cell differentiation. Indeed, blocking CD39 during *in vitro* T cell activation followed by *in vivo* transfer skewed differentiation towards an MPEC phenotype. CD39 and CD73 were further differentially expressed at a memory timepoint where we found increased CD39 expression on CD69+CD103+ resident memory T cells across several non-lymphoid tissues in mice. We further provide evidence to support these findings in humans, suggesting that T_RM_ express CD39. Finally, we found that virus-specific bystander T_RM_ in tumors can express comparable levels of CD39 to tumor-reactive CD8+ T cells.

In conclusion, this study is the first to demonstrate that CD39 and CD73 have distinct expression patterns on memory T cell precursors, and that modulating CD39 enzymatic activity during *in vitro* priming has effects on memory establishment. In addition, we have expanded upon our understanding of CD39 and CD73 on memory CD8+ T cells, illustrating distinct expression patterns on discrete memory subsets. This study has implications for our understanding of CD39 and CD73 as functional ecto-enzymes throughout T cell differentiation, which could lead to new strategies aimed at modulating T cell memory and implicate the purinergic signaling pathway in T_RM_ maintenance.

## Results

### The antiviral effector T cell response is associated with dynamic modulation of CD39 and CD73

It has been established that CD39 is expressed on exhausted CD8+ T cells responding to tumors^20,21,23,22^ and chronic viral infection^19^, but the kinetics of expression on antigen-specific CD8+ T cells *in vivo* during acute viral infection is unknown. To assess expression of purinergic ecto-enzymes on virus-specific T cells, we transferred 1 x 10^4^ naive transgenic OT-I CD8+ T cells specific for the model antigen ovalbumin into recipient C57BL/6 mice. The following day, mice were infected intranasally with vesicular stomatitis virus expressing ovalbumin (VSV_ova_), and sequential blood draws were taken on day 7, 10 and 14 post-infection (**Fig. 1a**). By day 7, there was an increased frequency of CD39+ OT-I T cells that remained elevated through day 14 compared to naive CD8+ T cells (**Fig. 1b**). In contrast, CD73 was decreased on OT-I cells compared to naive CD8+ T cells at day 7, but its expression increased over time (**Fig. 1c**). Importantly we saw the same expression patterns on CD44+ CD45.1-endogenous CD8+ T cells, strongly suggesting that this phenomenon is not specific to transgenic OT-I T cells **(Supplemental Fig. 1a-c**). While OT-I T cells are primarily CD39-CD73-on day 7 post infection, there was an increased diversity of CD39 and CD73 populations throughout viral clearance, with each subset represented about equally by day 14 (**Fig. 1d-e**). Thus, dynamic expression of CD39 and CD73 are associated with differentiation of antigen-specific CD8+ T cell following viral infection.

**Figure 1.**
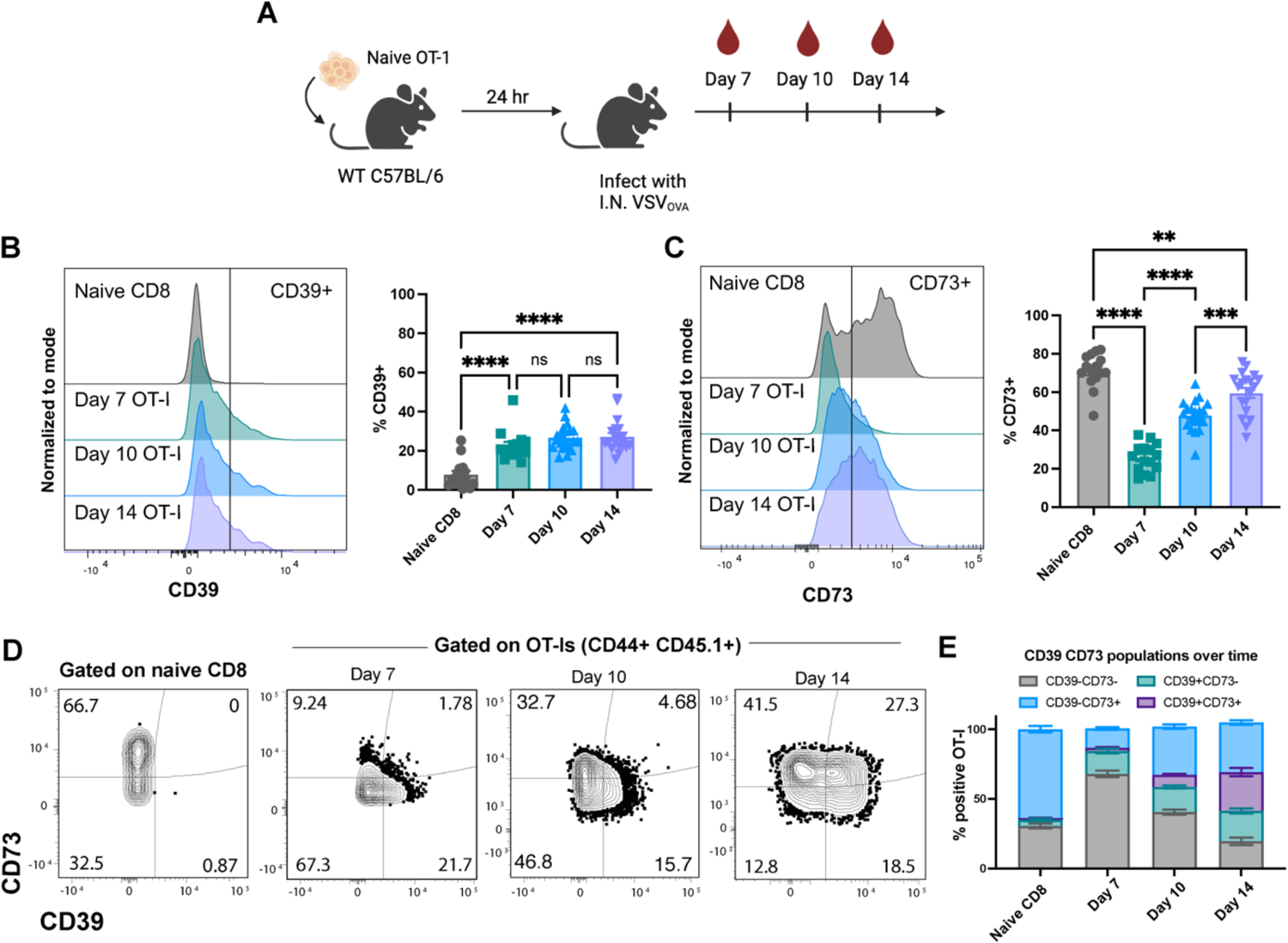
Anti-viral effector T cell response is associated with dynamic modulation of CD39 and CD73. **a.** Experimental schematic utilized to track effector virus-specific OT-I T cells in response to an acute VSV_ova_ infection. **b.** Representative histogram and combined bar graph depicting CD39 expression on OT-I T cells (CD44+ CD45.1+) and naive CD8+ T cells (day 7, CD44-CD45.1-) on day 7, 10, and 14 post infection in blood. **c.** Representative histograms and combined bar graph depicting CD73 expression on OT-I T cells (CD44+ CD45.1+) and naive CD8+ T cells (day 7, CD44-CD45.1-) on day 7, 10, and 14 post infection in blood. **d.** Representative flow plots showing CD39 and CD73 expression on naïve or OT-I T cells in blood, quantified over time in **e.** Data combined from two experiments. Naive CD8+ and day 7 OT-I n=15, day 10 and day 14 OT-I n=20. **b-c.** One-way ANOVA with Tukey’s post-test. *p <0.05, **p<0.01, ***p<0.001, ****p<0.0001, ns= not significant. Error bars represent the mean ± SEM.

### CD39 and CD73 are differentially expressed on effector T cell subsets

Given the importance of the extracellular ATP and ADO balance in modulating the T cell response, we next sought to understand whether CD39 and CD73 were associated with differentiation into memory precursor (MPEC) or short-lived effector cells (SLEC)^2,3,4^. We identified both MPECs (CD127+KLRG1-) and SLECs (CD127-KLRG1+) within circulating OT-I populations in the blood on day 7-14 (**Fig. 2a-b**). Strikingly, we found that CD39 expression was enriched in the SLEC population through all timepoints (**Fig. 2c-d**). We found the inverse pattern of expression when examining CD73 where there was an increased percentage of CD73+ MPECs compared to SLECs (**Fig. 2e-f**). Phenotyping the CD44+ CD45.1-endogenous CD8+ population yielded the same association of CD39 and CD73 with SLEC and MPECs respectively, suggesting that this is not an OT-I-dependent finding **(Supplemental Fig 1d-e**). Through co-expression analysis, we identified that a proportion of both MPECs and SLECs are CD39-CD73-, but SLECs are primarily CD39+CD73-, while MPECs are primarily CD39-CD73+ on day 7, 10 and 14 post-infection. By day 14, SLECs are comprised of a mix of all four populations, while MPECs are predominantly CD73+ CD39+/-. Consistent with figure 1e, the CD39-CD73-population in both MPECs and SLECs decreases over time. (**Fig. 2g-h**). Together, these results demonstrate that CD39 and CD73 are differentially expressed on memory precursor and short-lived effector T cell subsets.

**Figure 2.**
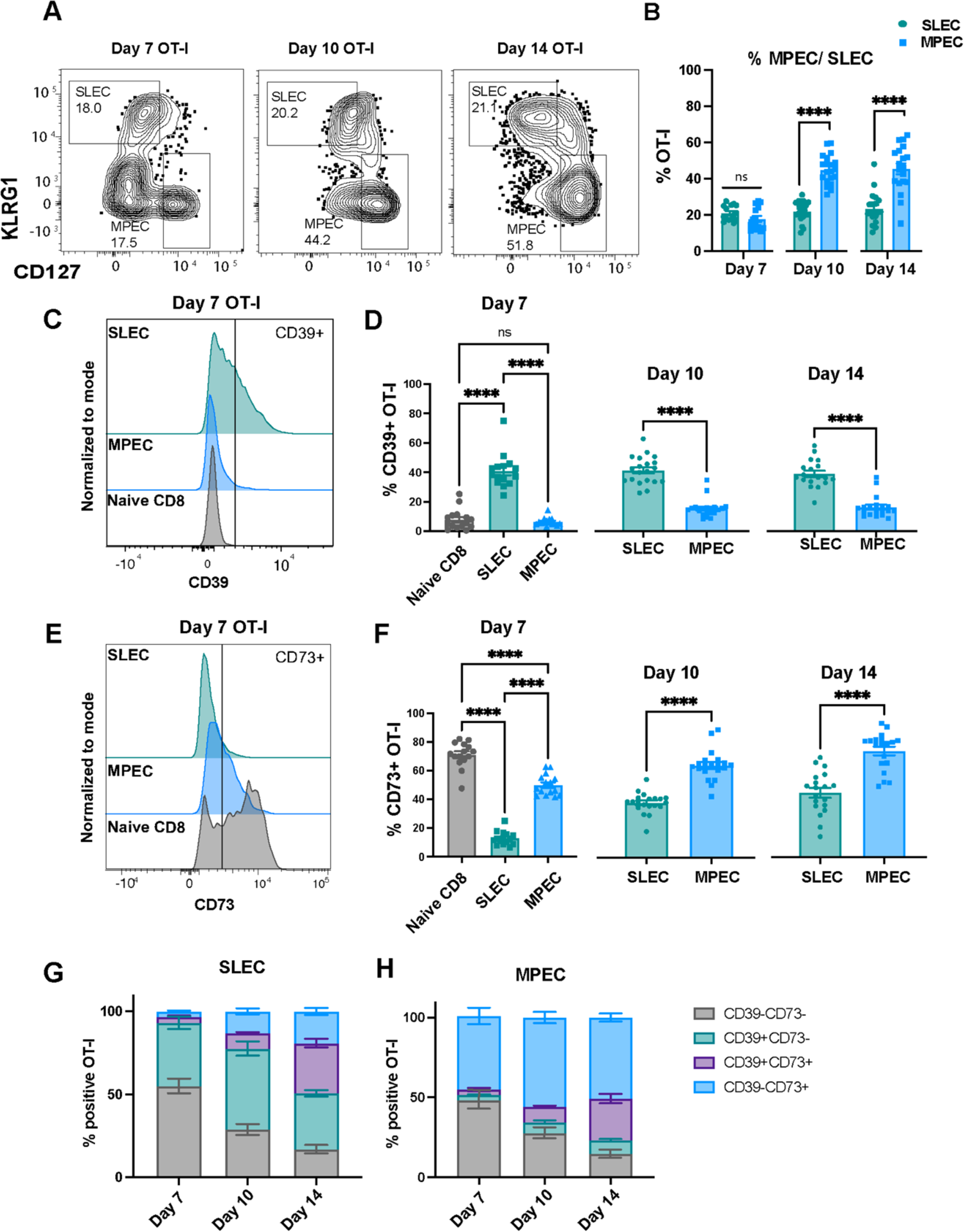
CD39 is increased on short-lived effector CD8+ T cells while CD73 is expressed on memory precursors during an acute anti-viral response. CD45.1+ OT-I were adoptively transferred into naive mice and infected with VSV_ova_ as in Fig 1a. **a.** Memory precursor; MPEC (CD127+KLRG1-) and short-lived effector; SLEC (CD127-KLRG1+) OT-I populations on day 7, 10, and 14 post infection in blood. **b.** Percent MPEC and SLEC of total OT-I T cells in blood over time. **c.** Representative histogram of CD39 expression on naive CD8+ T cells (CD44-CD45.1-) and day 7 MPEC or SLEC OT-I, quantified over time in **d. e.** Representative histogram of CD73 expression on naive CD8+ T cells (CD44-CD45.1-) and day 7 MPEC or SLEC OT-I, quantified over time in **f.** Vertical lines in **c** and **e** indicate gating for CD39+ and CD73+. **g-h.** Percent of CD39+/-CD73+/-OT-I T cells within the SLEC **(g)** and MPEC **(h)** compartment. Data combined from two experiments. Naive CD8+ and day 7 OT-I n=15, day 10 and day 14 OT-I n=20. **d** (day 7), **f** (day 7) One way ANOVA with Tukey’s post-test. **b, d, f.** Unpaired t test. *p <0.05, **p<0.01, ***p<0.001, ****p<0.0001, ns= not significant. Error bars represent the mean ± SEM.

### CD39 inhibition during in vitro T cell priming enhances memory establishment

Given the differential expression of CD39 on distinct memory precursor populations, we wanted to assess if CD39 enzymatic activity may be functionally skewing CD8+ T cell differentiation. To do this, we employed an *in vitro* priming model where we utilized a small molecule inhibitor (POM-1) to block CD39 activity during T cell activation (**Fig 3a**). In brief, we cultured splenocytes from an OT-I mouse for three days with SIINFEKL peptide and IL-2, with or without POM-1. The following day, we transferred 1 x 10^6^ activated OT-I T cells from each condition into separate naive recipient mice and infected with VSV_ova_ the following day (**Fig. 3a**). We found that *in vitro* priming in the presence of the CD39 blockade significantly enhanced OT-I frequency in the blood by day 14 (**Fig 3d**). In addition, the POM-1 treated OT-I T cells had an enhanced percentage of MPECs and decreased percentage of SLEC at day 7 and day 14 (**Fig. 3b-c, e-f).** The MPEC skewing with CD39 blockade was not seen *in vitro* before *in vivo* transfer (not shown). Interestingly, POM-1 treatment did not alter CD39 or CD73 on circulating OT-I *in vivo*, suggesting that differences in ecto-enzyme expression did not drive the observed phenotype **(Supplemental Fig. 2a-b**). At memory (>30 days post infection), there was a trending increase in POM-1-treated circulating OT-I T cells in the blood, LN, and spleen, though not statistically significant (**Fig. 3i**). We next assessed establishment of resident memory T cells. To ensure we were limiting our analyses to OT-I T cells within the tissue parenchyma, we utilized intravenous labeling to discriminate cells within the vasculature from resident in tissue^34,35^. While resident memory establishment was highly variable in this model, there was a statistically significant increase in the frequency of POM-1 treated OT-I in the brain and salivary gland with a similar trend across other tissues (**Fig. 3g-h, j).** There was also a statistically significant increase in total number of POM-1 treated OT-I in the brain and small intestine compared to untreated OT-I **(Supplemental Fig. 2e-k**). Additionally, we did not find any differences in CD69 or CD103 expression among OT-I T_RM_ across non-lymphoid tissues between experimental groups **(Supplemental Fig. 2l).** As seen in the circulating compartment (**Supplemental Fig. 2a-b**), there was also generally no difference in CD39 or CD73 expression on OT-I in tissues between the two experimental groups at memory **(Supplemental Fig. 2c-d**). In sum, CD39 inhibition during *in vitro* T cell priming enhances memory precursors in circulation and resident memory establishment in select non-lymphoid tissues.

**Figure 3.**
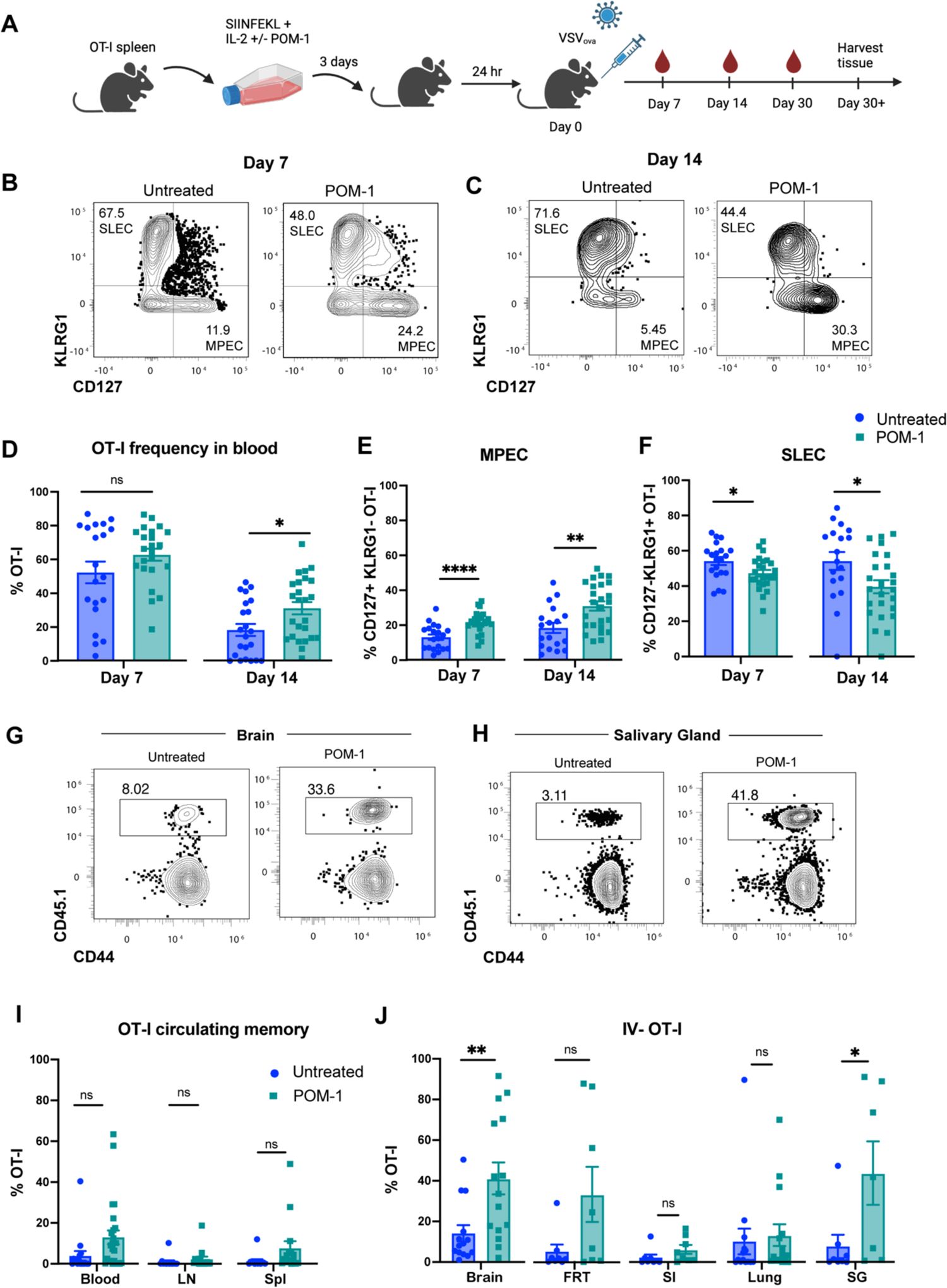
Inhibiting CD39 during T cell in vitro priming leads to OT-I T cell MPEC skewing and enhanced T_RM_ establishment. **a.** Experimental schematic demonstrating *in vitro* OT-I T cell priming in the presence or absence (+/-) of POM-1 (50uM), followed by transfer and subsequent infection *in vivo*. **b-c.** Representative flow plots showing CD127 and KLRG1 staining in untreated and POM-1 treated OT-I T cells on day 7 **(b)** and day 14 **(c)** in blood. **d.** Frequency of OT-I T cells in blood on day 7 and 14 post infection. **e-f**. Frequency of MPEC **(e)** and SLEC **(f)** populations among OT-I T cells primed +/-POM-1 in blood. **g-h.** Representative flow plots of CD45.1+ OT-I populations in brain **(g)** and SG **(h)** within each experimental group. **i.** Frequency of total OT-I T cells found in blood, lymph node or spleen 30+ days post infection. **j.** Frequency of IV-negative OT-I T cells in tissues 30+ days post infection (gated on IV-negative CD45.1 or CD90.1) FRT= female reproductive tract, SG= salivary gland, SI= small intestine, LN= lymph node, ns= not significant. Data in B-F combined from 3 independent experiments. Untreated n=20, POM-1 n=25. Data in G from three experiments (blood n=17 untreated, n=25 POM-1). Data in H from 2-3 experiments (brain and lung from 3 experiments, untreated n= 14, POM-1 n= 15; FRT, SI, Lung, SG from 2 experiments, n=8 per group). Unpaired t test. *p <0.05, **p<0.01, ***p<0.001, ****p<0.0001, ns= not significant. Error bars represent the mean ± SEM.

### CD39 and CD73 are co-expressed on resident memory T cells

Given the variation in CD39 and CD73 expression during the effector phase, we next sought to characterize expression at a memory timepoint. To do this, we employed the same mouse model as in Fig. 1a and analyzed resident and circulating OT-I T cells 40-50 days after VSV_ova_ infection, which has been previously shown to establish T_RM_ across tissues^36^. We surveyed CD39 and CD73 co-expression on circulating and resident T cells from the lymph node and non-lymphoid tissues and found distinct expression patterns (**Fig. 4a**). As before, when characterizing T_RM_, we utilized intravenous labeling to limit our analysis to cells within tissue excluded from the vasculature. We found that CD73 was ubiquitously expressed on all memory T cell subsets, while CD39 expression varied significantly (**Fig. 4b**). The CD69+ CD103+ T_RM_ had a significantly increased frequency of CD39+ cells followed by T_EM_ and T_CM_ (**Fig. 4c**). Further, co-expression of CD39 and CD73 was found at the highest frequency in CD69+CD103+ T_RM_, while CD39-CD73+ OT-I were predominantly found in T_CM_ (**Fig. 4d**). The tissues we analyzed demonstrated a diversity of CD69 and CD103 expression on IV-negative OT-I T cells (**Fig. 4e-f**). When IV-negative T cells were subset based on CD69 and CD103 expression, we found a remarkably consistent expression pattern of CD39 and CD73 across T_RM_ in every non-lymphoid tissue analyzed. Interestingly, the frequency of CD39+CD73+ OT-I increased from CD69-CD103- to CD69+CD103- and CD69+CD103+ across every NLT analyzed, with over 75% of CD69+CD103+ T_RM_ co-expressing CD39 and CD73 in each tissue (**Fig. 4g**). In contrast, CD39+CD73-OT-I T cells were associated with CD69+CD103-T_RM_ (**Fig. 4h**). Moreover, CD39-CD73+ OT-I were more enriched in the CD69-CD103-population (**Fig. 4i**). In sum, CD39+CD73+ co-expression is associated with tissue residency.

**Figure 4.**
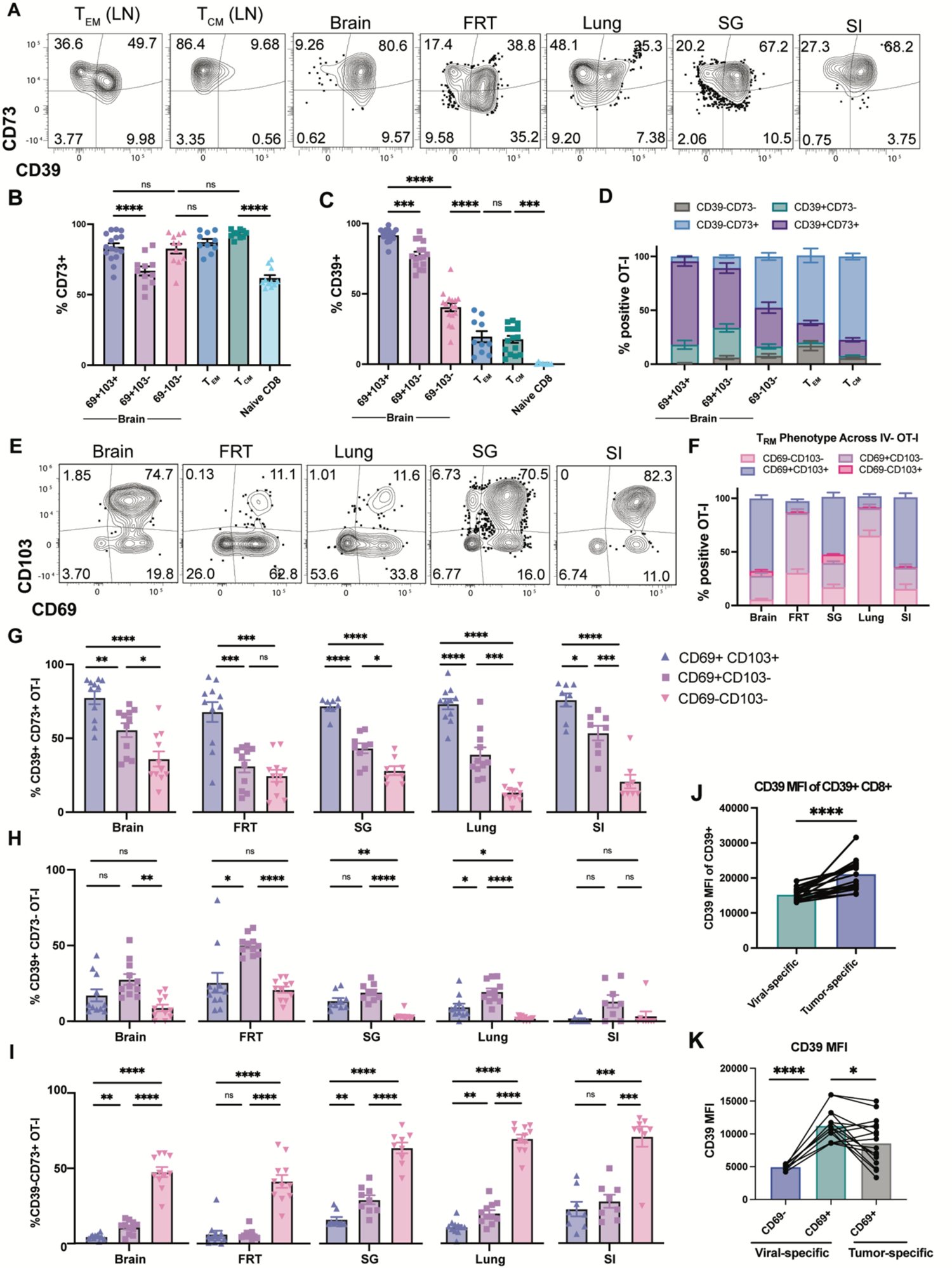
CD39 and CD73 are expressed on T_RM_. Memory OT-I T cells were harvested from mice at day 40-50 post infection and CD39/CD73 expression was analyzed across memory subsets. **a**. Flow plots depicting CD39 and CD73 expression on effector memory (T_EM_) and central memory (T_CM_) OT-I from the superficial cervical lymph node and IV-negative OT-I across non-lymphoid tissues. **b-c.** Frequency of CD73+ **(b)** or CD39+ **(c)** OT-I T_RM_ (CD69+/CD103+/-) in the brain, OT-I T_EM_ (CD62L-), and OT-I T_CM_ (CD62L+) in the lymph node and naïve (CD44-) CD8+ T cells in spleen. **d.** Frequency of CD39+/-CD73+/- IV-negative OT-I T_RM_ in the brain, and OT-I T_EM_ and T_CM_ in the lymph node. **e.** Representative flow plots of CD69 and CD103 residency phenotype across non-lymphoid tissues, quantified in **f. g-i.** Frequency of CD39+CD73-OT-I **(g)**, CD39+CD73-OT-I **(h)** and CD39-CD73+ OT-I **(i)** stratified by CD69 and CD103 expression across tissue. **j-k**. Mice were infected intravenously with vaccinia virus, then received an adoptive transfer of 1 x 10^6^ *in vitro* activated OT-I T cells 30 days later. 7 days after transfer, mice received B16-ova brain tumors and were euthanized at day 16 post tumor challenge. Vaccinia-specific endogenous CD8+ T cells were identified by B8R H2-K^b^ tetramer. **j.** MFI of CD39, gated on IV-negative CD39+ OT-I T cells (tumor-specific) or IV-negative B8R+ T cells (virus-specific). Paired data are shown where each line connects datapoints from individual mice. **k.** MFI of CD39 on CD69+ tumor-specific OT-I T cells, and CD69+ and CD69-memory virus-specific B8R+ T cells. Gated on IV-negative. T_EM_= CD44+ CD45.1+ CD69-CD103-CD62L-, T_CM_= CD44+ CD45.1+(or Thy1.1+) CD69-CD103-CD62L+, T_RM_= IV-CD44+CD45.1+(or Thy1.1+) CD69+CD103+/-. FRT= female reproductive tract, SG= salivary gland, SI= small intestine, LN= lymph node. **a-i.** Data combined from 2-3 experiments. n= 10-16 mice. **j-k.** Two independent experiments, n=17 mice. (b-c, g-i, k) One way ANOVA, (j) unpaired t test. *p <0.05, **p<0.01, ***p<0.001, ****p<0.0001, ns= not significant. Error bars represent the mean ± SEM.

### CD39 inhibition does not affect T_RM_ ex vivo recall response

Given that CD39+ CD8+ T cells in tumors and chronic viral infection express phenotypic markers of exhaustion and have decreased polyfunctionality^19,20^, we wanted to understand how CD39 expression on T_RM_ impacts their function. To test this, we generated OT-I memory mice as in Fig 1a., then harvested T cells from brain and spleen at a memory timepoint and cultured them *ex vivo* **(Supplemental Fig. 3).** The majority of T cells from the brain are IV-negative and CD69+CD103+/-thus express CD39, while spleen OT-I are comprised of circulating populations and are CD39^lo^ (**Fig. 4**). We pre-treated the isolated brain T_RM_ or spleen OT-I for one hour with or without POM-1, then reactivated with control irrelevant (gp33) or cognate (SIINFEKL) peptide for 4 hours and measured cytokine production. As previously demonstrated^8,37^, OT-I T cells from the brain were functional and upregulated IFNγ and TNFα in response to cognate peptide **(Supplemental Fig. 3a, c, e)**. We observed no significant differences in production of IFNγ or TNFα when cells were reactivated in the presence of POM-1 **(Supplemental Fig. 3c).** Together, while CD39 may have significant effects on CD8+ T cell differentiation during priming, CD39 activity does not appear to modulate *ex vivo* T_RM_ reactivation.

### CD39 is expressed at similar levels on CD69+ CD103+ bystander T_RM_ and tumor-reactive TILs in mice

Given the recent findings that cancer unrelated (bystander) TILs do not express CD39^20^ and that CD39+CD103+ CD8+ T cells are tumor-reactive^22^, we sought to fit our data with these existing paradigms. To directly compare CD39 expression on tumor and bystander virus-specific CD8+ T cells, we utilized a B16-ova brain metastasis model. We first established a virus-specific memory T cell population by infecting mice with vaccinia virus. Over 30 days later, we transferred *in vitro* activated OT-I T cells, and 7 days later challenged mice with B16-ova in the dorsal striatum of the brain **(Supplemental Fig. 4a).** Sixteen days post tumor inoculation, we assessed IV-negative CD8+ T cells using H-2Kb tetramer to identify anti-viral T cells specific for the immunodominant epitope of vaccinia virus (B8R) or tumor-specific OT-I T cells (**Supplemental Fig. 4b,c)**. In agreement with previous reports^20,22,24–27,28^, total tumor-specific CD8+ T cells had increased CD39 expression compared to total virus-specific T cells (**Fig 4j**). However, when cells were subset by CD69 expression, CD69+ virus-specific T cells expressed increased levels of CD39 compared to tumor-specific T cells **(Supplemental Fig. 4b-c**, **Fig 4k)**. These findings suggest that bystander CD69+ CD8+ T cells in tumors can express CD39 at similar levels to tumor-specific T cells in mice. Thus, CD39 may be associated with a tissue residency phenotype independent of chronic antigen stimulation.

### Evidence that CD39 is expressed on T_RM_ in humans

We next wanted to investigate whether our observations about CD39 and residency translated to humans. To do this, we first procured autopsy tissues from four deceased male and female individuals from 36-76 years of age (see methods). We identified T_RM_ subsets, defined by CD69 and CD103 expression, on polyclonal CD45RA-CCR7-memory CD8+ T cells in brain tissue, including cortex and meninges, as well as in lung and skin (**Fig 5a-b**). While not statistically significant, we found increased CD39 expression on CD69+CD103+ T_RM_ compared to CD69-CD103-T cells across tissues (**Fig. 5c-f**). Expanding on our mouse tumor findings, (**Fig. 4j-k)** we next analyzed Epstein Barr Virus (EBV)-specific CD8+ T cells identified by HLA-A*02 combinatorial tetramer staining in a human melanoma tumor^38^. As seen previously, EBV-specific T cells in the tumor expressed CD69 and CD103 (Fig. 5g)^29,39^. We found that over 70% of both CD69+CD103+ EBV-specific T_RM_ and non-viral specific T_RM_ expressed CD39 to similar levels (**Fig. 5h)**. These data suggest that CD39 can be expressed on CD69+ CD103+ T_RM_ in human tissues, including bystander T_RM_ in melanoma tumors.

**Figure 5.**
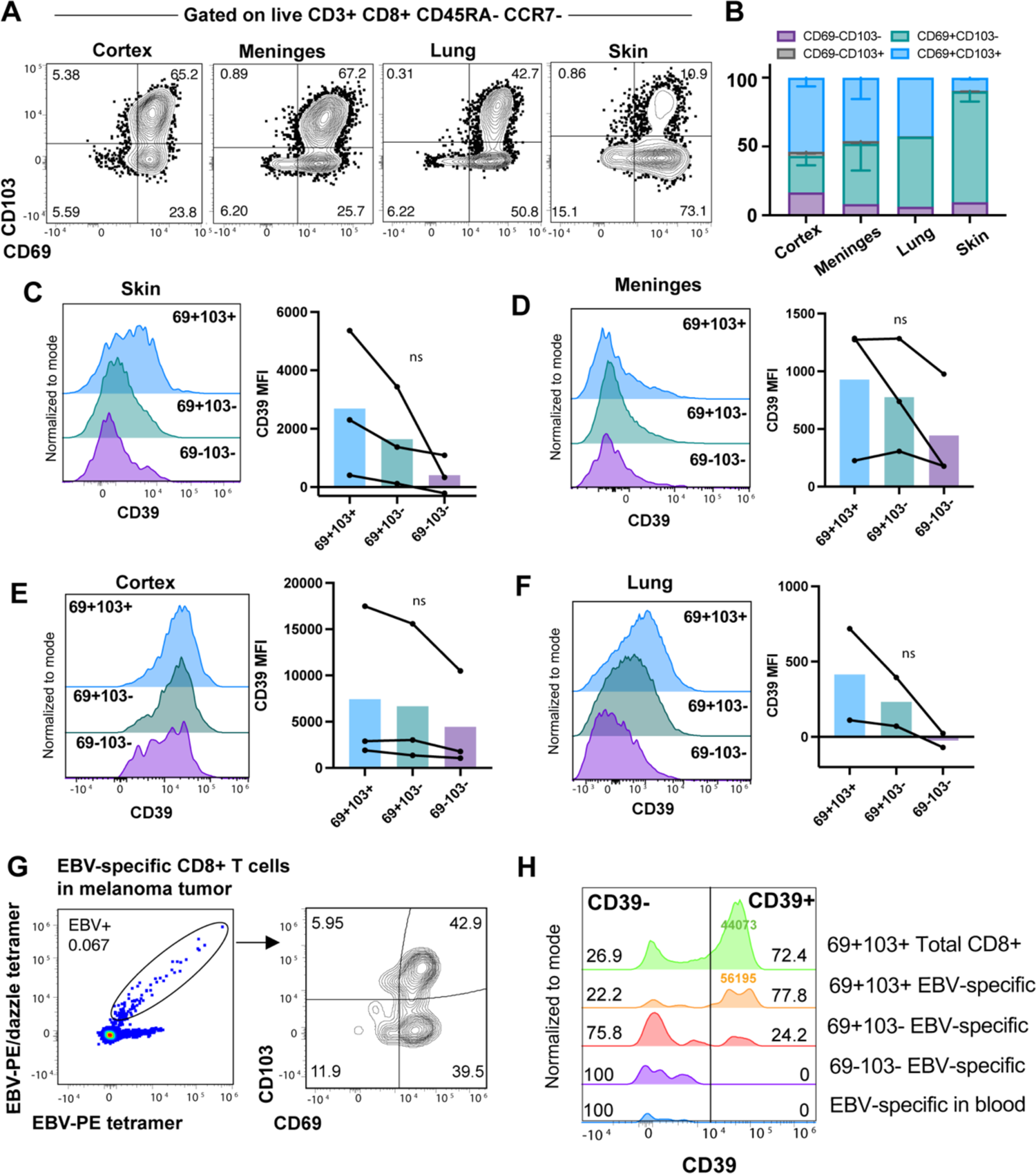
CD39 is expressed on CD69+ CD103+/-CD8+ T cells in human tissue. **a.** Representative flow plots showing CD69 and CD103 expression on CD45RA-CCR7-memory CD8+ T cells in human autopsy tissues quantified in **(b). c-f.** Representative histograms of CD39 expression on CD8+ T cells. CD39 MFI across CD69+/CD103+, CD69+/CD103-, and CD69-/CD103-CD8+ T cells, quantified in bar graphs. Graphs connecting individual patient data for skin **(c),** meninges **(d)**, cortex **(e)** and lung **(f).** Data points from three individual decedents connected via line plot. **g.** Identification of Epstein Barr Virus (EBV)-specific CD8+ T cells in a patient melanoma tumor by combinatorial HLA-A2 tetramer staining (cells gated on live, CD8+ CD3+ CD45RA-CCR7-). Flow plot depicting CD69 and CD103 expression on EBV-specific cells (right). **h.** CD39 expression on total CD45RA-CCR7-EBV_GLC_-specific CD8+ T cells in blood (blue) compared to EBV-specific CD69+/-CD103+/-T cells in the tumor (purple, red, orange), compared to CD45RA-CCR7-non-viral specific CD8+ T cells in the tumor (green). All samples gated on live, CD8+ CD3+ CD45RA-CCR7-. Numbers in black are percentages, colored numbers inset in peaks are MFI. Autopsy samples n=4, patient data in supplemental table 1. Melanoma patient sample= 1. **c-f.** Paired one way ANOVA. Error bars represent the mean ± SEM.

## Discussion

Our work here shows that CD39 and CD73 are dynamically expressed on T cells over the course of differentiation following an acute viral infection. This study extends the role of CD39 beyond a marker of tumor-specific or exhausted CD8+ T cells, and highlights its role during T cell differentiation and memory. We found that CD39 expression increased on T cells following infection, consistent with previous *in vitro* and *in vivo* studies^18,40^. Additionally, we found a unique association of each ecto-enzyme with distinct effector subsets; CD39 was expressed predominantly on short-lived effector cells, while CD73 was restricted to memory precursor cells. Interestingly, it’s been shown that activated human CD39+ CD4+ T cells *in vitro* are more apoptotic than CD39-CD4+ T cells due to activation of the AMPK-p53 pathway^40^. Pharmacological inhibition and transfection of CD39 dampened and enhanced this phenotype, respectively, illustrating a direct functional role of CD39 in apoptosis^40^. Our data expand on these findings by demonstrating that CD39 is expressed predominantly on CD8+ SLECs, which are prone to apoptosis and do not develop into long-lived memory cells^41^. Thus, an attractive hypothesis given this study and our data is that CD39 expression on SLECs may contribute to their ultimate cell death during the contraction phase of T cell differentiation.

In contrast to SLECs, we found that CD73 was expressed by MPECs, which go on to form long-lived memory. We also identified CD73 on all memory subsets including circulating and resident across several non-lymphoid tissues. The role of CD73 on memory precursors and differentiated memory subsets is likely through the production of adenosine (ADO). Signaling through the adenosine receptor (A_2A_R) can prevent TCR-induced downregulation of CD127 (IL-7R), which is found on memory precursors and can promote their survival^31,42^. Thus, CD73 on memory T cells could serve as a continuous source of adenosine, which signals through A_2A_R and supports survival^43,44,45^. Furthermore, human circulating memory CD73+ CD4+ T cells have increased expression of pro-survival genes as well as reduced propensity to undergo apoptosis, further emphasizing the functional role of CD73 in memory T cell survival^46^.

Functionally, we found that CD39 inhibition during *in vitro* T cell priming skews T cell differentiation away from short-lived effectors and towards memory precursors. Given the role of CD39 in apoptosis and T cell contraction, inhibition of CD39 ATPase activity during *in vitro* T cell priming may prevent these apoptotic signals. CD39 inhibition could also be skewing T cell differentiation through modulation of T cell receptor (TCR) signaling strength, which dictates precursor establishment^30^. TCR signal strength is directly related to eATP signaling through P2X7R and subsequent Ca^2+^ release, thus perturbing the eATP balance through CD39 blockade *in vitro* may modulate this pathway^47^. However, in our *in vitro* priming studies, we cannot distinguish between T cell intrinsic or extrinsic effects as our culture contained a mixed population of T cells, B cells and antigen-presenting cells (APCs), which can all express various levels of CD39^48,49^. Thus, the CD39 inhibitor could be blocking ATPase activity on other cell types, resulting in a general increase in ATP levels in the culture. In sum, perturbing the eATP balance through CD39 blockade *in vitro* may shift the proportions of precursor cells through TCR signaling strength, however, further studies are required to elucidate this mechanism.

While CD73 was expressed on all memory subsets, CD39+ CD73+ cells were enriched in CD69+ CD103+ T_RM_. Tissue resident virus-specific CD8+ T cells have been previously shown to differentially express CD39 and CD73^33^. However, the relative expression of CD39 and CD73 on circulating and resident memory subsets, as well as within the resident compartment had not been characterized prior to this study. While CD73 was expressed on all memory subsets, we identified that CD39 was found at highest frequencies on T_RM_ in mice and present evidence that this trend is consistent in humans. Further, we discovered distinct co-expression patterns on T cell subsets, where T_EM_ and T_CM_ were generally CD39-CD73+ and T_RM_ were positive for both ecto-enzymes. This suggests a novel finding of purinergic ecto-enzymes in T cell biology; that memory CD8+ T cells generally express CD73 and that CD39 expression is dictated by residency.

In addition to distinct ecto-enzyme expression patterns across memory subsets, we identified significant differences in subsets within the T_RM_ compartment. The fact that CD39 and CD73 are only co-expressed on CD69+CD103+ T_RM_ is quite striking. Ecto-enzyme co-expression could suggest an enhanced dependence on purinergic signaling associated with tissue residency, collaborating in tandem to reduce eATP and increase ADO levels. The low-affinity ATP receptor, P2X7R, can induce cell death in CD8+ T cells depending on its relative expression^50,51^. Indeed, high levels of P2X7R on CD8+ T_RM_ can trigger cell death during tissue damage allowing for new antigen-specific cells to take residence^14,50,51^. Thus, CD39 and CD73 co-expression on T_RM_ could be a potential mechanism to divert excess ATP from the extracellular space, preventing unnecessary P2X7R-mediated cell death and promoting T_RM_ homeostatic survival. In parallel, the increased ADO production could enhance survival through A_2A_R^42^. To further this point, we found that *ex vivo* reactivation of brain OT-I T_RM_ was not affected by CD39 blockade, which expands upon other studies with CD73 blockade^33^. This suggests that CD39 and CD73 expression may not control the function of T_RM_ but could be working in tandem to promote survival through modulation of ATP/ADO.

CD39, often with co-expression of CD103, has been described as a marker of tumor-specific T cells ^19,20,21,22,23^. To align our data with these studies, we assessed whether CD39 was restricted to tumor-specific CD8+ T cells compared to bystander virus-specific T_RM_. When CD39 was analyzed agnostically to T_RM_ phenotype, CD39 expression was increased on tumor-specific CD8+ T cells compared to virus-specific, which is consistent with previous reports^20,22^. We were surprised to find that when stratified by CD69 expression, tumor-specific CD8+ T cells did not express increased CD39 compared to virus-specific CD69+ T_RM_. Consistent with this, we found that EBV-specific CD69+CD103+ T_RM_ in a human melanoma tumor expressed similar levels of CD39 as total CD8+ T cells which likely includes any tumor-specific TILs. However, there are limitations to our studies. While we identified EBV-specific T cells by tetramer, we did not include tetramers to detect tumor-specific populations, rather we analyzed total memory T cells which include mixed specificities. Additionally, while our mouse model enables the study of bystander and tumor-specific T cells in parallel, our analysis is limited to CD69 as a T_RM_ marker since this tumor model does not induce CD103 expression (data not shown). In all, our studies show that CD39 is expressed by bystander T_RM_ in tumors and suggests that CD39 should be used cautiously as an absolute marker of tumor-specific TILs.

Here, we demonstrated that CD39 and CD73 are associated with distinct short-lived effector and memory precursors, and that CD39 inhibition modulates the differentiation trajectory of CD8+ T cells. These data open the opportunity to explore CD39 inhibitors during *in vitro* activation as a potential method of modulating memory precursors and T_RM_ establishment in a clinical setting, such as adoptive cell therapies. Beyond the classical understanding of CD39 and CD8+ T cell exhaustion, our results demonstrate that CD39 and CD73 are expressed during viral clearance and define distinct memory subsets. Co-expression of CD39 and CD73 is associated with CD69+CD103+ T_RM,_ identifying a novel pathway involved in the putative maintenance of T_RM_. In all, our data suggest that the CD39/CD73 axis is an important regulator of CD8+ T cell differentiation and implicate the purinergic signaling pathway in T_RM_ maintenance.

## Acknowledgements

We thank the Rosato and Skorput labs for their intellectual input, moral support, and collaboration. We thank the DartLab Immune Monitoring Core for maintaining the flow cytometers and the Dartmouth Animal Care Facility Staff members for skilled animal husbandry. We thank Dr. Steven Fiering and Dr. Patricia Pioli for providing manuscript direction and thesis advisory. Experiment schematics made with BioRender.com. We thank David Masopust for providing VSV_ova_ and VV_gp33_ viruses. This work was supported by bioMT through NIH NIGMS grant P20-GM113132, NCI R01 CA269455-01A1, and NCI Cancer Center Support Grant 5P30CA023108-37.

## Article contributions

JFI and PCR conceptualized the project and designed the experiments. JFI organized and conducted the experiments. HND optimized and processed several of the human autopsy and melanoma experiments with technical and intellectual support from TGS. MJT provided tumor lines, human melanoma tissue and intellectual input. SAK, TC, SCM, and MAF helped with mouse tissue processing. CL and GJZ procured autopsy tissues and provided feedback on tissue processing optimization. AGJS helped with statistical analysis and gave intellectual input. JFI and PCR interpreted the results, wrote, and edited the manuscript. All authors approved the submitted manuscript.

## Materials and Methods

### Mice

6-10 week female C57BL/6J mice were purchased from the Jackson Laboratory (Bar Harbor, ME) and housed under specific pathogen free conditions in the Dartmouth College vivarium. Experimental mice were used at 6-14 weeks of age. CD45.1+ and CD90.1+ OT-I mice were fully backcrossed to C57BL/6J mice and bred in our animal colony. All studies were approved by the Institutional Animal Care and Use Committee at Dartmouth College.

### Adoptive transfers and infections

OT-I memory mice were generated by transferring 1 x 10^4^ naive CD90.1+ or CD45.1+ CD8+ OT-I T cells from female mice into naive female CD45.2+ C57BL/6J recipients, then infecting with 1 x 10^6^ PFU vesicular stomatitis virus expressing ovalbumin (VSV_ova_) i.n. after 24 hours. Mice were screened in advance of experiments for circulating CD44+ CD45.1+ (or CD90.1+) OT-I populations to assess memory establishment. For B16-ova brain tumor studies, CD45.2+ C57BL/6 mice were infected with 1 x 10^7^ PFU/mL vaccinia virus expressing gp33 (VV_gp33_)^52^ i.v. and received 1 x 10^6^ *in vitro* activated CD45.1+ OT-I cells 30 days post infection. Vaccinia virus-specific CD8+ T cells were stained with H-2K^b^/B8R MHC I tetramer (made in-house as previously described)^53^.

### Lymphocyte isolation from tissues

We used a previously described intravascular staining method to exclude cells in the vasculature from cells in tissue^34,35^. We injected mice i.v. with biotin or Alexa Fluor 488 conjugated anti-CD8α and euthanized the mice 3 minutes later for tissue processing as described^35^. Tissues were stained with CD39-perCP-eF710 (Clone 24DMS1, BioLegend 5015962, 1:100), CD73-FITC (Clone Ty/11.8, BioLegend 127220, 1:100), CD8b-BV711 (Clone SK1, BioLegend 126633, 1:1000), Streptavidin-BV650 (BioLegend 405232, 1:250), CD45.1-BUV737 (Clone A20, BD 612811, 1:250), CD44-BV785 (Clone IM7, BioLegend 103006, 1:500), CD69-PE-dazzle (Clone H1.2F3, BioLegend 104536, 1:75), CD103-APC (Clone 2E7, BioLegend 121414, 1:100), PD-1-PE-Cy7 (Clone 29F.1A12, BioLegend 135216, 1:50), KLRG1-PE-dazzle (Clone 2F1/KLRG1, BioLegend 138424, 1:100), CD127-PE-Cy5 (Clone SB/199, BioLegend 121123, 1:200), CD62L-BV510 (Clone MEL-14, BioLegend 104441, 1:500), and Ghost Dye Red 780 (Cytek 12-0365-T100, 1:25,000). Cells were stained for 30 minutes at 4 degrees and analyzed on an Aurora spectral cytometer (Cytek).

B16-ova tumors were isolated from tissue, cut into small pieces, and digested in 0.5mg/ml type IV collagenase for 30 minutes at 37 degrees while spinning. Tumors were then mechanically disrupted by gentleMACS dissociator (Miltenyi) (setting m_spleen) and lymphocytes were separated using a 44/67% Percoll density gradient (Cytvia). B16-ova tumors were i.v labeled with CD8α-Alexa 488 (Clone 53-6.7, BioLegend 100723). Cells were blocked for 10 minutes with Fc block (DartLab Immune Monitoring Core), then surface stained with CD39-perCP-eF710 (Clone 24DMS1, BioLegend 5015962, 1:100), CD8b-BV711 (Clone SK1, BioLegend 126633, 1:1000), CD4-BV421 (Clone GK1.5, BioLegend 100438, 1:500), CD45.1-BUV737 (Clone A20, BD 612811, 1:250), CD44-BV785 (Clone IM7, BioLegend 103006, 1:500), CD69-PE-dazzle (Clone H1.2F3, BioLegend 104536, 1:75), CD103-BV510 (Clone 2E7, BioLegend 121423, 1:100), PD-1-PE-Cy7 (Clone 29F.1A12, BioLegend 135216, 1:50), CD62L-Alexa 700 (Clone MEL-14, BioLegend 104426, 1:200), H2-K^b^ B8R-PE tetramer (made in house, 1:50) and Ghost Dye Red 780 (Cytek 12-0365-T100, 1:25,000). Samples were analyzed on the Aurora Spectral Cytometer (Cytek).

### Blood processing and staining

Blood was obtained through submandibular bleeds into heparin-coated Eppendorf tubes, then transferred to 5mL FACS tubes. Red blood cells were lysed with ACK lysis buffer for 2 minutes, centrifuged at 1600rpm for 5 minutes, lysed with ACK for 2 minutes, then diluted in tissue media or FACS buffer followed by a final centrifugation at the above settings. Blood was stained with CD39-perCP-eF710 (Clone 24DMS1, BioLegend 5015962, 1:100), CD73-FITC (Clone Ty/11.8, BioLegend 127220, 1:100), CD8b-Alexa 647 (Clone YTS1567.7, BioLegend 126611, 1:100), CD45.1-BUV737 (Clone A20, BD 612811, 1:100), CD44-BV785 (Clone IM7, BioLegend 103006, 1:500), PD-1-PE-Cy7 (Clone 29F.1A12, BioLegend 135216, 1:50), KLRG1-PE-dazzle (Clone 2F1/KLRG1, BioLegend 138424, 1:100), CD127-PE-Cy5 (Clone SB/199, BioLegend 121123, 1:200), CD62L-BV510 (Clone MEL-14, BioLegend 104441, 1:500), and Ghost Dye Red 780 (Cytek 12-0365-T100, 1:25,000). Cells were stained for 30 minutes at 4 degrees and analyzed on an Aurora Spectral Cytometer (Cytek).

### Intracranial B16-ova tumors

B16-ova tumor lines were generated in-house and generously shared by the Turk Laboratory (Dartmouth)^54^. Mice were anesthetized and given 3.25 mg/kg slow-release buprenorphine (Ethiqua XR) s.c. prior to tumor implantation. Following sanitation of the surgical area with sterile 70% ethanol wipes, an incision was made on the midline to expose the skull. The mouse was positioned in a Stereotactic Injector (Stoelting 53311) and a burr hole was made to the left of the midline at 1mm lateral and 1mm posterior from bregma using a micromotor high-speed drill (Stoelting 51449). Next, a 10ul Hamilton syringe was placed above the burr hole and descended 4mm posterior then 2mm dorsal to create a 2mm pocket. 3ul of 150,000 B16-ova cells was injected into the dorsal striatum at 1ul/minute, followed by a 3-minute rest before removal of the syringe. Povidone iodine was dabbed on the surface of the burr hole to prevent tumor growth out of the skull. Bone wax (SMI Z046) was used to cover the burr hole and the incision was closed with Vetbond tissue adhesive (3M 1469SB). Mice were allowed to recover on a heating pad and monitored daily until the 16-day experiment endpoint.

### In vitro T cell priming and blockade with POM-1

OT-I spleens were processed as above and cultured in T175 flasks with T cell media (500mL RPMI 1640 + 2.0mM L-glutamine, 50 mL heat inactivated FBS, 5ml 1 M HEPES, 5 ml 100x non-essential amino acids, 5ml 100x sodium pyruvate, 1.75ul 2-ME). Cells were treated with POM-1 (Tocris cat #2689, 50uM), recombinant human IL-2 (R&D systems, cat # 202-IL-010/CF), and SIINFEKL peptide (0.5ug, Biomatik) for 3 days. ‘Untreated’ activated OT-I T cells were cultured with SIINFEKL and IL-2. OT-I T cells were observed morphologically for signs of T cell activation (clustering) and CD8 T cell frequency and activation phenotype was analyzed by flow cytometry. Mice received 1 x 10^6^ POM-1 or untreated OT-I T cells on day-1, followed by intranasal infection with VSV_ova_ the following day.

### Ex vivo T cell reactivation

Spleen and brain were harvested from 3-5 OT-I memory mice and processed as described in ‘lymphocyte processing from tissues’ section. Samples were pooled and plated in triplicate per condition in a 96-well flat bottom plate with T cell media. POM-1 (50uM) was added to samples and incubated for 1 hour. Control peptide (gp33, 0.5ug, Biomatik), reactivating peptide (SIINFEKL, 0.5ug, Biomatik), or 12-myristate 13-acetate (PMA) and ionomycin (1:500) was added to the cells for 4-6 hours in the presence of Invitrogen™ eBioscience™ Protein Transport Inhibitor Cocktail (Cat #50-112-9032)After 4-6 hours, the cells were harvested and stained with CD39-perCP-eF710 (Clone 24DMS1, BioLegend 5015962, 1:100), CD8b-Alexa 647 (Clone YTS1567.7, BioLegend 126611, 1:20,000), CD45.1-BUV737 (Clone A20, BD 612811, 1:100), CD44-BV785 (Clone IM7, BioLegend 103006, 1:500), CD69-PE-dazzle (Clone H1.2F3, BioLegend 104536, 1:75), CD103-BV510 (Clone 2E7, BioLegend 121423, 1:100), Ghost Dye Red 780 (Cytek 12-0365-T100, 1:25,000). Cells were fixed and permeabilized using the Foxp3 Transcription Factor Fix/Perm Concentrate (Cytek, TNB-1020-L050) and Diluent (Cytek, TNB-1022-L160). Cells were then intracellularly stained with IFNγ-BV650 (Clone XMG1.2, BioLegend 505832, 1:100) and TNFα-BV421 (Clone MP6-XT22, BioLegend 506327).

### Human autopsy and tumor samples

Human autopsy samples were procured from the Department of Pathology at Dartmouth Health (Lebanon, NH). Tissues were obtained from four decedents (50% male, 50% female) between ages 36 and 76 who passed from pneumonia (n=2), myocardial infarction, and *Capnocytophaga canimorsus* infection. Cases were selected for inclusion in the study based on time processed, where each sample was obtained within 24 hours of death to ensure viability of cells for flow cytometry. Lung was excluded from autopsies who passed of pneumonia. The melanoma tumor sample and paired blood were obtained from a standard-of-care resection at Dartmouth Health according to IRB guidelines (Lebanon, NH). Blood samples were diluted in PBS and overlayed onto 4mL Ficoll (Cytvia) in a 15mL tube, then spun for 15 minutes at 400xg room temperature. The PBMC layer was collected into a 15mL tube and diluted in RPMI, then spun at 300xg for 8 minutes. Cells were rinsed with FACS buffer (DPBS 1x, 1g Bovine Serum Albumin, 5mL 10% Na Azide) and stained with the same panel below. Lymph nodes, cortex, lung, skin and tumor were cut into small pieces, and digested in 0.5mg/ml (cortex) or 1mg/ml (lung, skin, meninges, tumor) type IV collagenase plus DNAse (1:500) for 30 minutes (cortex), 1 hour (meninges), or 2 hours (lung, skin, tumor) at 37 degrees while spinning. Tissues were then mechanically disrupted by gentleMACS dissociator (setting m_spleen) and lymphocytes were separated using a 44/67% Percoll density gradient. Autopsy tissues were stained with LIVE/DEAD Fixable Blue Dead Cell Stain Kit, for UV excitation (Thermo Fisher L23105, 1:500) for 30 minutes at 4 degrees Celsius. Cells were then washed and surface stained with CD4-PE-Cy5 (Clone RPA-T4, BioLegend 300510, 1:400), CD69-BUV496 (Clone FN50, BD 750214, 1:30), CD39-BUV737 (Clone TU66, BD 612852, 1:30), CD45RA-BV650 (Clone HI100, BioLegend 304136, 1:400), CD103-BV711 (Clone Ber-ACT8, BioLegend 350222, 1:100), PD-1-PE-Cy7(Clone EH12.2H7, BioLegend 329918, 1:30), CD3-Alexa Fluor 700 (Clone UCHT1, BioLegend 300424, 1:100), CD8-APC-Fire 750 (Clone SK1, BioLegend 344746, 1:400). Following viability stain, the melanoma tumor was then stained with HLA-A2*01-restricted virus-specific tetramers (in house, 1:100) loaded with EBVBML(GLCTLVAML) and EBVLMP(CLGGLLTMV) peptides 1:100 for 30min at 4Cto detect CD8+ T cells specific for Epstein Barr Virus (EBV). Cells were subsequently surface stained using the panel above and data was acquired using the Aurora Spectral Cytometer (Cytek).

### Cell Definitions

Naïve CD8+ T cell: CD8+ T cell that has not encountered antigen and is CD44-(mouse). Memory CD8+ T cell: CD8+ T cells that have encountered antigen (when known) >30 days ago and/or are CD44+ (mouse), or CCR7-CD45RA-(human). Memory precursor effector cell: CD8+ T cells that have encountered antigen <30 days ago and are CD44+ CD127+ KLRG1-. Short-lived effector cell: CD8+ T cells that have encountered antigen <30 days ago and are CD44+ CD127-KLRG1+. Effector memory T cell (T_EM_): CD8+ T cells that have encountered antigen >30 days ago and are CD44+ CD62L-CD69-CD103-. Central memory T cell (T_CM_): CD8+ T cells that have encountered antigen >30 days ago and are CD44+ CD62L+ CD69-CD103-. Resident memory T cell (T_RM_): CD8+ T cells that have encountered antigen >30 days ago and are IV-CD44+ CD69+ CD103+/-.

### Statistical analysis

Statistical analysis was performed using GraphPad Prism. Data was subjected to the Shapiro-Wilk normality test to determine whether they were sampled from a Gaussian distribution. All datasets demonstrated Gaussian distribution. One-way ANOVA tests were performed for multiple comparisons with Tukey post-test. Paired or unpaired Student’s t test were completed for two comparisons. Data are represented as mean + SEM. P values < 0.05 were considered significant. All experiments were analyzed using Flowjo 10.10 and Prism 10 (GraphPad).

## Supplementary Materials

This article includes Supplemental Figs. 1-4.

## References

1. Wherry, E. J. & Ahmed, R. Memory CD8 T-Cell Differentation during Viral Infection. J. Virol. 78, 5535–5545 (2004).

2. Cui, W. & Kaech, S. M. Generation of effector CD8+ T cells and their conversion to memory T cells. Immunol. Rev. 236, 151–166 (2010).

3. Molecular regulation of effector and memory T cell differentation | Nature Immunology. www.nature.com/articles/ni.3031.

4. Huster, K. M. et al. Selective expression of IL-7 receptor on memory T cells identifies early CD40L-dependent Generation of distinct CD8+ memory T cell subsets. Proc. Natl. Acad. Sci. U. S. A. 101, 5610–5615 (2004).

5. Jameson, S. C. & Masopust, D. Understanding subset diversity in T cell memory. Immunity 48, 214–226 (2018).

6. Rosato, P. C., Beura, L. K. & Masopust, D. Tissue resident memory T cells and viral immunity. Curr. Opin. Virol. 22, 44–50 (2017).

7. Mueller, S. N. & Mackay, L. K. Tissue-resident memory T cells: Local specialists in immune defence. Nat. Rev. Immunol. 16, 79–89 (2016).

8. Schenkel, J. M. & Masopust, D. Tissue-resident memory T cells. Immunity 41, 886–897 (2014).

9. Smolders, J. et al. Tissue-resident memory T cells populate the human brain. Nat. Commun. 9, 1–14 (2018).

10. Behr, F. M. et al. Tissue-resident memory CD8+ T cells shape local and systemic secondary T cell responses. Nat. Immunol. 21, 1070–1081 (2020).

11. Szabo, P. A., Miron, M. & Farber, D. L. Location, location, location: Tissue resident memory T cells in mice and humans. Sci. Immunol. 4, 1–12 (2019).

12. Vardam-Kaur, T. et al. Pannexin-1 channels promote CD8+ T cell effector and memory responses through distinct intracellular pathways. bioRxiv 2023.04.19.537580 (2023) doi:10.1101/2023.04.19.537580.

13. Wanhainen, K. M., Jameson, S. C. & da Silva, H. B. Self-Regulation of Memory CD8 T Cell Metabolism through Extracellular ATP Signaling. Immunometabolism 1, e190009 (2019).

14. Vardam-Kaur, T. et al. The Extracellular ATP Receptor P2RX7 Imprints a Promemory Transcriptional Signature in Effector CD8+ T Cells. J. Immunol. Author Choice 208, 1686–1699 (2022).

15. Bono, M. R., Fernández, D., Flores-Santibáñez, F., RosemblaX, M. & Sauma, D. CD73 and CD39 ectonucleotidases in T cell differentation: Beyond immunosuppression. FEBS LeE. 589, 3454–3460 (2015).

16. Timperi, E. & Barnaba, V. CD39 Regulation and Functions in T Cells. Int. J. Mol. Sci. 22, 8068 (2021).

17. Antonioli, L., Pacher, P., Vizi, E. S. & Haskó, G. CD39 and CD73 in immunity and inflammation. Trends Mol. Med. 19, 355–367 (2013).

18. Raczkowski, F. et al. CD39 is upregulated during activation of mouse and human T cells and aXenuates the immune response to Listeria monocytogenes. PLOS ONE 13, e0197151 (2018).

19. Gupta, P. K. et al. CD39 Expression Identifies Terminally Exhausted CD8+ T Cells. PLoS Pathog. 11, 1–21 (2015).

20. Simoni, Y. et al. Bystander CD8+ T cells are abundant and phenotypically distinct in human tumour infiltrates. Nature 557, 575–579 (2018).

21. Lee, Y. J., et al. CD39 + tissue-resident memory CD8 + T cells with a clonal overlap across compartments mediate antitumor immunity in breast cancer. Sci. Immunol. 7, eabn8390 (2022).

22. Duhen, T. et al. Co-expression of CD39 and CD103 identifies tumor-reactive CD8 T cells in human solid tumors. Nat. Commun. 9, (2018).

23. Canale, F. P. et al. CD39 expression defines cell exhaustion in tumor-infiltrating CD8+ T cells. Cancer Res. 78, 115–128 (2018).

24. Eiva, M. A., Omran, D. K., Chacon, J. A. & Powell, D. J. Systematic analysis of CD39, CD103, CD137, and PD-1 as biomarkers for naturally occurring tumor antigen-specific TILs. Eur. J. Immunol. 52, 96–108 (2022).

25. Goncharov, M. M. et al. Pinpointing the tumor-specific T cells via TCR clusters. eLife 11, e77274 (2022).

26. Zhu, W. et al. CD8+CD39+ T Cells Mediate Anti-Tumor Cytotoxicity in Bladder Cancer. OncoTargets Ther. 14, 2149–2161 (2021).

27. Fehlings, M. et al. Checkpoint blockade immunotherapy reshapes the high-dimensional phenotypic heterogeneity of murine intratumoural neoantigen-specific CD8+ T cells. Nat. Commun. 8, (2017).

28. Chow, A. et al. The Ectonucleotidase CD39 Identifies Tumor-reactive CD8+ T cells Predictive of Immune Checkpoint Blockade Efficacy in Human Lung Cancer. Immunity 56, 93–106.e6 (2023).

29. Rosato, P. C. et al. Virus-specific memory T cells populate tumors and can be repurposed for tumor immunotherapy. Nat. Commun. 10, (2019).

30. Knudson, K. M., Hamilton, S. E., Daniels, M. A., Jameson, S. C. & Teixeiro, E. Cuqng Edge: The Signals for the Generation of T Cell Memory Are Qualitatively Different Depending on TCR Ligand Strength. J. Immunol. 191, 5797–5801 (2013).

31. Cekic, C., Sag, D., Day, Y.-J. & Linden, J. Extracellular adenosine regulates naive T cell development and peripheral maintenance. J. Exp. Med. 210, 2693–2706 (2013).

32. Crompton, J. G. et al. Akt inhibition enhances expansion of potent tumor-specific lymphocytes with memory cell characteristics. Cancer Res. 75, 296–305 (2015).

33. Smith, C. J. & Snyder, C. M. Inhibitory Molecules PD-1, CD73 and CD39 Are Expressed by CD8+ T Cells in a Tissue-Dependent Manner and Can Inhibit T Cell Responses to Stimulation. Front. Immunol. 12, (2021).

34. Anderson, K. G. et al. Intravascular staining for discrimination of vascular and tissue leukocytes. Nat. Protoc. 9, 209–222 (2014).

35. Steinert, E. M. et al. Quantifying Memory CD8 T Cells Reveals Regionalization of Immunosurveillance. Cell 161, 737–749 (2015).

36. Nelson, C. E. et al. Robust Iterative Stimulation with Self-antigens Overcomes CD8+ T Cell Tolerance to Self- and Tumor antigens. Cell Rep. 28, 3092–3104.e5 (2019).

37. Schøller, A. S., Nazerai, L., Christensen, J. P. & Thomsen, A. R. Functionally Competent, PD-1+ CD8+ Trm Cells Populate the Brain Following Local antigen Encounter. Front. Immunol. 11, (2021).

38. Newell, E. W., Klein, L. O., Yu, W. & Davis, M. M. Simultaneous detection of many T-cell specificities using combinatorial tetramer staining. Nat. Methods 6, 497–499 (2009).

39. Ning, J. et al. Functional virus - specific memory T cells survey glioblastoma. Cancer Immunol. Immunother. (2021) doi:10.1007/s00262-021-03125-w.

40. Fang, F. et al. Expression of CD39 on Activated T Cells Impairs their Survival in Older Individuals. Cell Rep. 14, 1218–1231 (2016).

41. S S., et al. Functional and genomic profiling of effector CD8 T cell subsets with distinct memory fates. J. Exp. Med. 205, (2008).

42. Cekic, C. & Linden, J. Adenosine A2A receptors intrinsically regulate CD8+ T cells in the tumor microenvironment. Cancer Res. 74, 7239–7249 (2014).

43. RosemblaX, M. V. et al. Ecto-5ʹ-Nucleotidase (CD73) Regulates the Survival of CD8+ T Cells. Front. Cell Dev. Biol. 9, 647058 (2021).

44. Himer, L. et al. Adenosine A2A receptor activation protects CD4+ T lymphocytes against activation-induced cell death. FASEB J. 24, 2631–2640 (2010).

45. Hirata, Y. et al. CD150high Bone Marrow Tregs Maintain Hematopoietic Stem Cell Quiescence and Immune Privilege via Adenosine. Cell Stem Cell 22, 445–453.e5 (2018).

46. Fang, F. et al. The cell-surface 5ʹ-nucleotidase CD73 defines a Functional T memory cell subset that declines with age. Cell Rep. 37, 109981 (2021).

47. Yip, L. et al. Autocrine regulation of T-cell activation by ATP release and P2X7 receptors. FASEB J. 23, 1685–1693 (2009).

48. Zhao, H., Bo, C., Kang, Y. & Li, H. What Else Can CD39 Tell Us? Front. Immunol. 8, (2017).

49. Allard, B., Longhi, M. S., Robson, S. C. & Stagg, J. The ectonucleotidases CD39 and CD73: Novel checkpoint inhibitor targets. Immunol. Rev. 276, 121–144 (2017).

50. Aswad, F. & Dennert, G. P2X7 receptor expression levels determine lethal effects of a purine based danger signal in T lymphocytes. Cell. Immunol. 243, 58–65 (2006).

51. Stark, R. et al. TRM maintenance is regulated by tissue damage via P2RX7. Sci. Immunol. 3, eaau1022 (2018).

52. Schenkel, J. M. et al. Resident memory CD8 T cells trigger protective innate and adaptive immune responses. Science 346, 98–101 (2014).

53. Fitzpatrick, K. S. et al. Validation of Ligand Tetramers for the detection of antigen-Specific Lymphocytes. J. Immunol. 210, 1156–1165 (2023).

54. Byrne, K. T. et al. Autoimmune melanocyte destruction is required for robust CD8+ memory T cell responses to mouse melanoma. J. Clin. Invest. 121, 1797–1809 (2011).

