## Supplemental Figures 1-4 for "CD39 is expressed on functional effector and tissue resident memory CD8+ T cells"

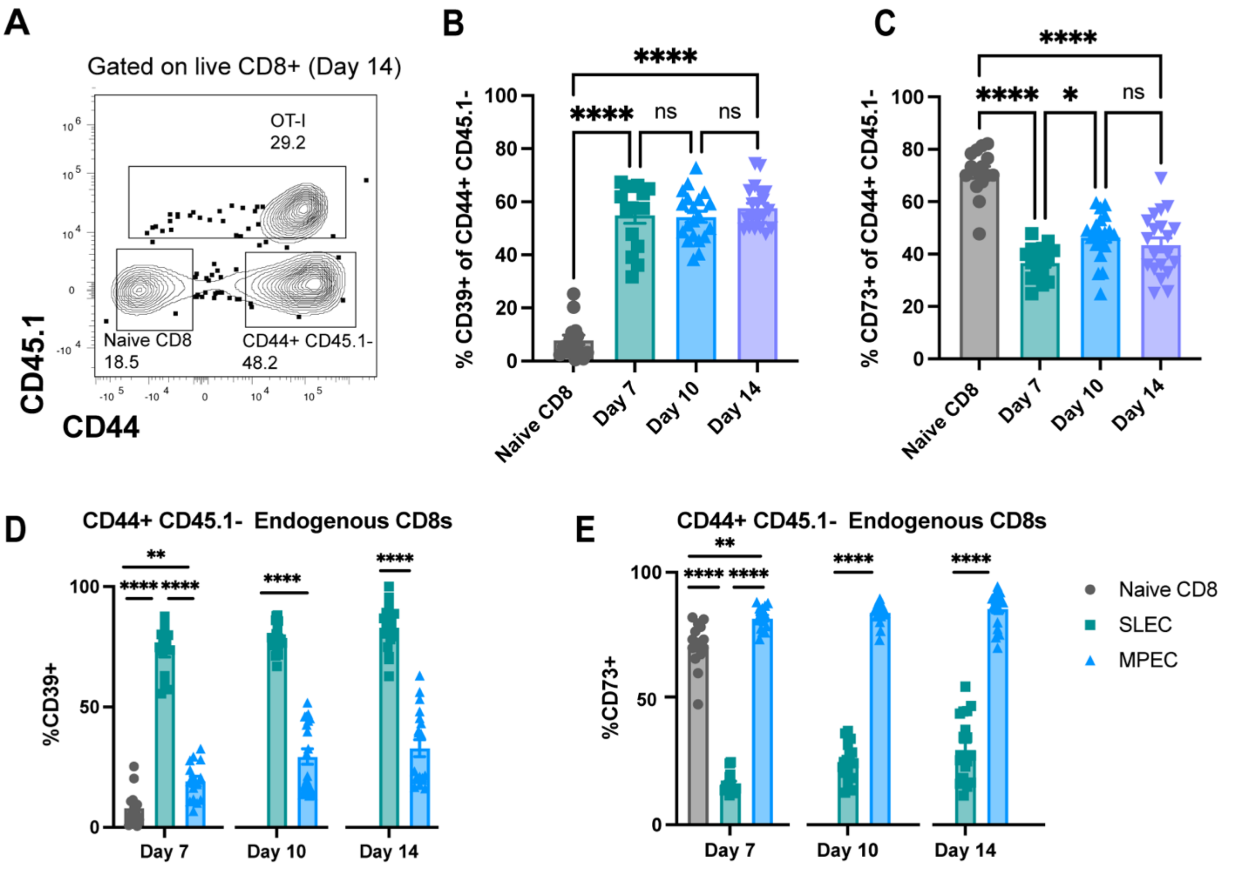


**Supplemental Figure 1. CD39 and CD73 expression on endogenous CD44+ CD45.1- CD8+ T cells after viral infection. a**. Identification of naïve CD8+ T cells, CD44+CD45.1- endogenous CD8+ T cells, CD44+CD45.1+ OT-I T cells by gating on live CD8+. **b.** Percent CD39+ CD8+ T cells within the endogenous CD44+ CD8+ population. **c.** Percent CD73+ CD8+ T cells within the endogenous CD44+ CD8+ population. **d**. Percent CD39+CD44+ CD8 T cells within SLEC (CD127-KLRG1+) and MPEC (CD127+KLRG1-) populations. **e.** Percent CD73+CD44+ CD8 T cells within SLEC (CD127-KLRG1+) and MPEC (CD127+KLRG1-) populations. Data combined from two independent experiments. Naive CD8+ and day 7 OT-I n=15, day 10 and day 14 OT-I n=20. **d** (day 7), **e** (day 7) One way ANOVA with Tukey's post test. **b, c.** Unpaired t test. *p <0.05, **p<0.01, ***p<0.001, ****p<0.0001, ns= not significant. Error bars represent the mean ± SEM.

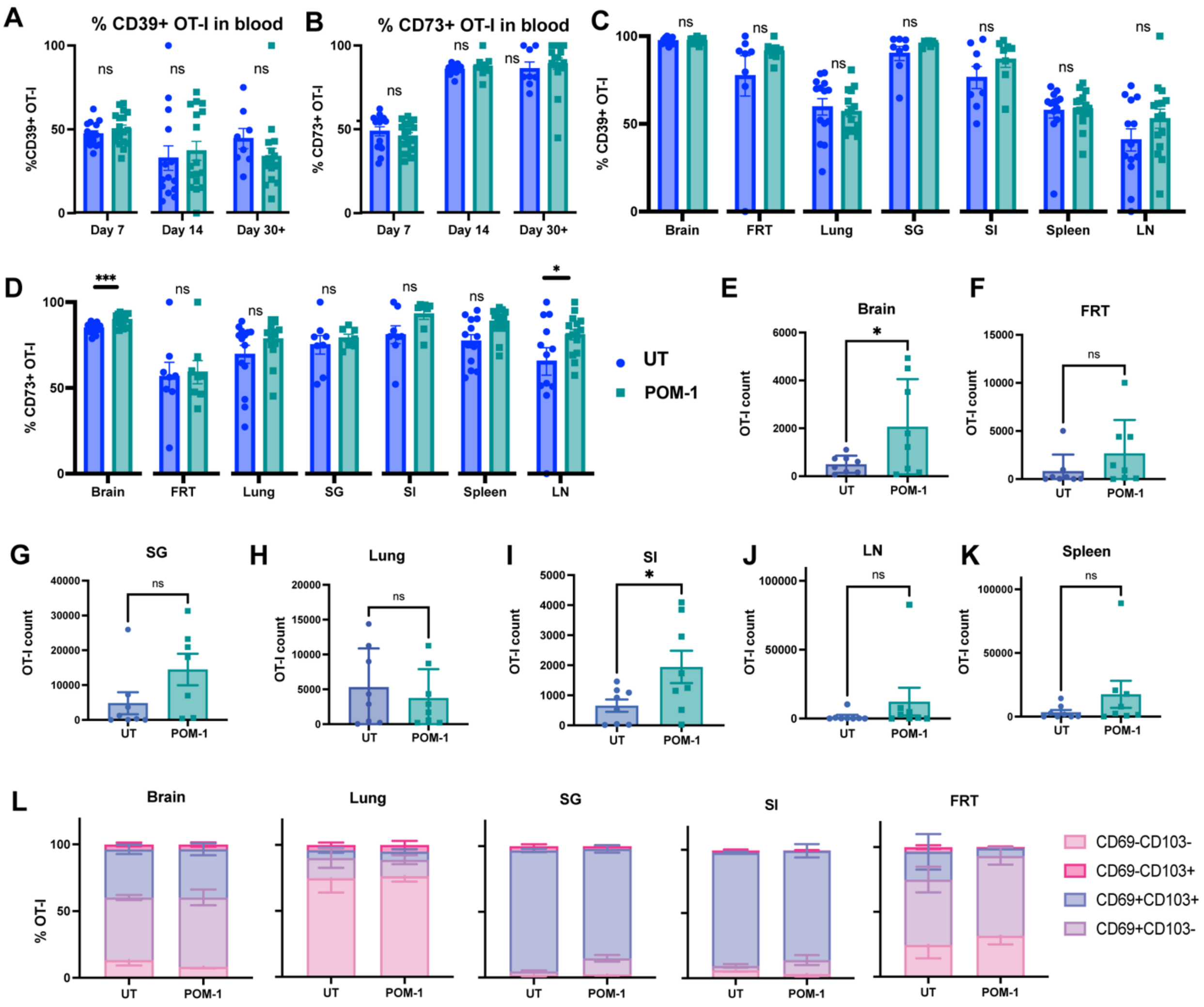


**Supplemental Figure 2. Effects of CD39 inhibition during *in vitro* T cell priming.** Experimental schematic outlined in figure 3a, mice euthanized at day 40-50 post infection. **a-b.** %CD39+ **(a)** and %CD73+ **(b)** OT-Is in blood across POM-1 and untreated groups from day 7-14. **c-d.** %CD39+ **(c)** and %CD73+ **(d)** OT-Is across tissues at day 30+. **e-k.** Total IV- OT-I T cell count in each tissue. **l.** CD69 and CD103 expression on OT-I T cells from non-lymphoid tissues in both groups. Non-lymphoid tissues and LN were gated on iv- cells, spleen gated on total CD8. UT= untreated, FRT= female reproductive tract, SG= salivary gland, SI= small intestine, LN= lymph node. Data combined from 2-3 independent experiments. n=8 POM-1, n=8 untreated, **a-k.** Unpaired student's t test. *p< 0.05, **p<0.01, ***p<0.001, ****p<0.0001, ns= not significant. Error bars represent the mean ± SEM.

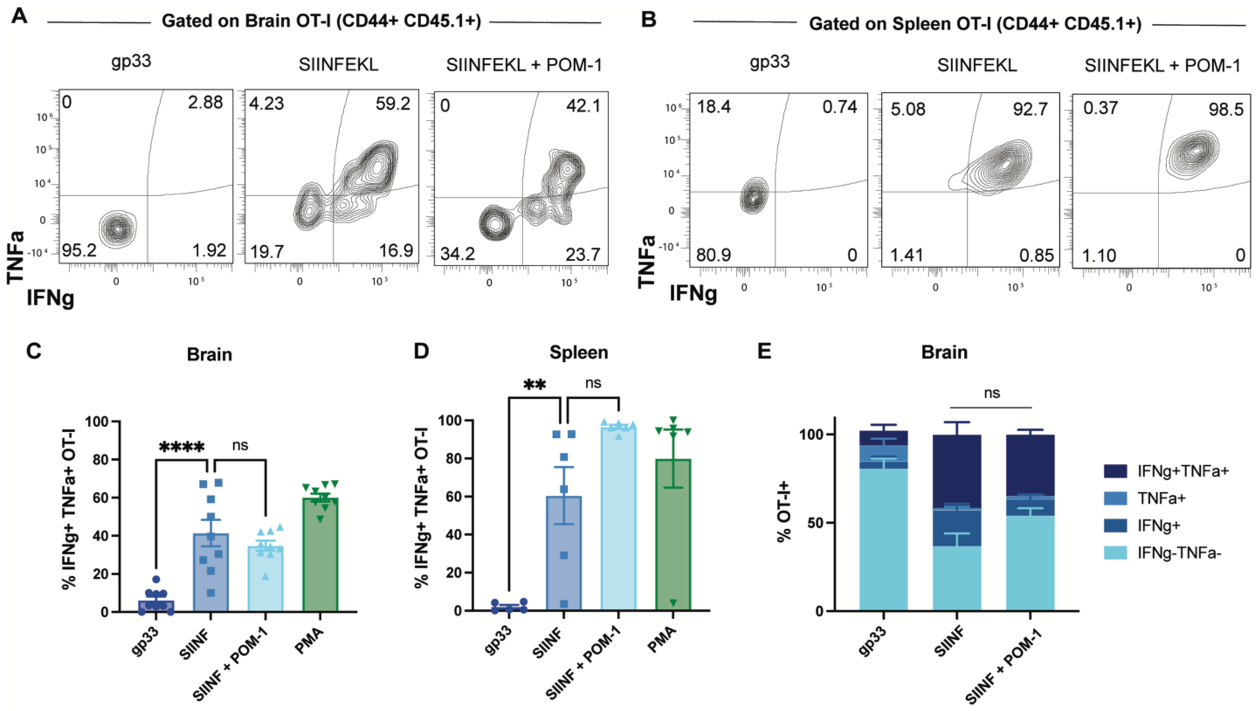


**Supplemental Figure 3. CD39 blockade does not affect *ex vivo* brain $T_{RM}$ reactivation.** Brain and spleen were harvested from OT-I memory mice as described in figure 1. Brain and spleen were pre-incubated for 1 hour with or without POM-1, then 4 hours with control (gp33) or viral (SIINFEKL) peptide. **a-b.** IFNγ and TNFα production by brain **(a)** and spleen **(b)** OT-I when treated with control peptide (gp33), cognate peptide (SIINFEKL), or cognate peptide with CD39 blockade (SIINFEKL + POM-1), quantified by percent positive in **c** and **d.** **e.** Percent IFNγ +/- TNFα +/- OT-I from brain in each experimental condition. Data combined from 2-3 experiments (brain 3 repeats, spleen 2 repeats), n=3 per group per experiment. One way ANOVA with Tukey's post-test. *p <0.05, **p<0.01, ***p<0.001, ****p<0.0001, ns= not significant. Error bars represent the mean ± SEM.

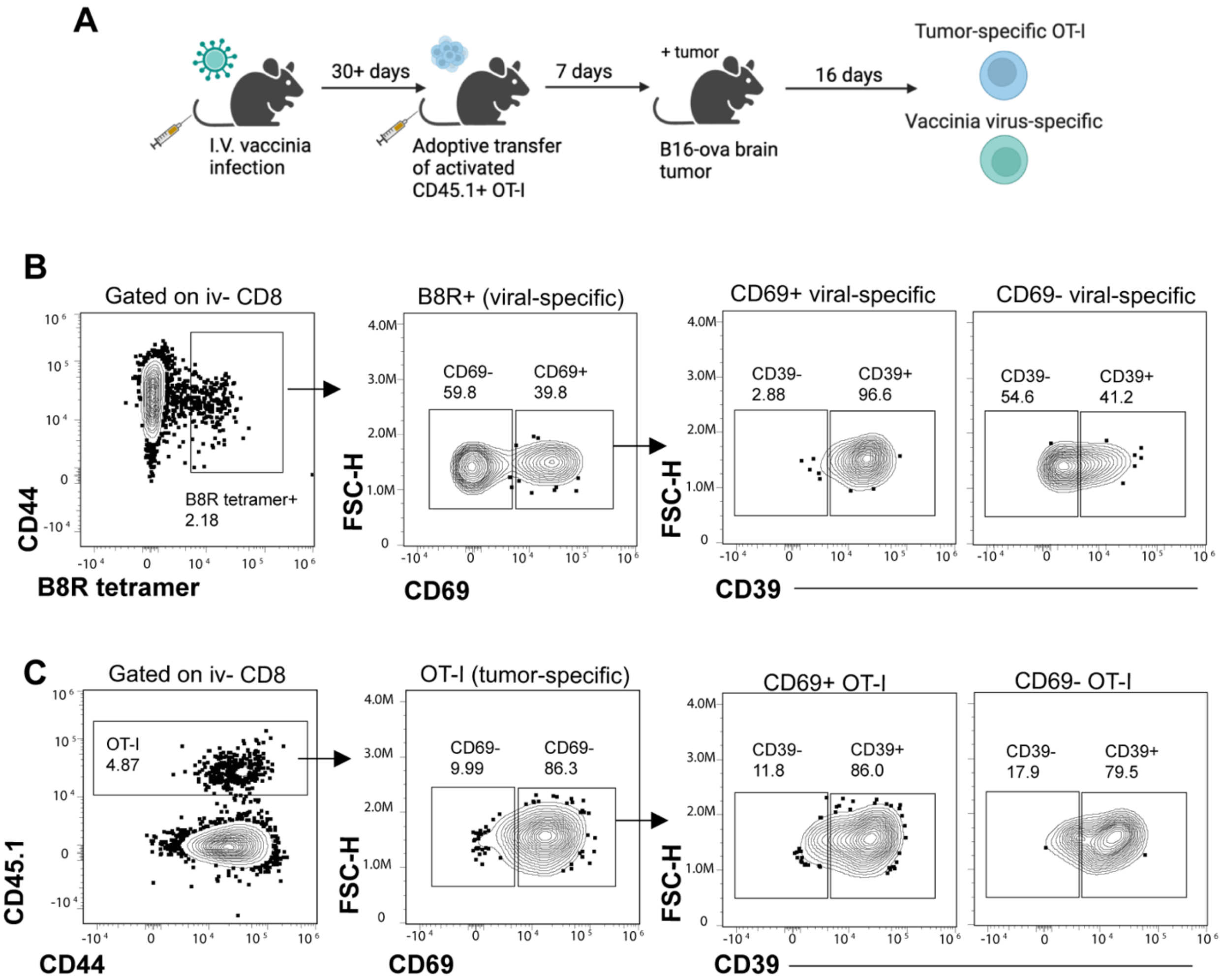


**Supplemental Figure 4. CD39 is expressed on CD69+ bystander $T_{RM}$ and tumor-reactive cells. a.** Experimental schematic used to generate memory vaccinia virus-specific CD8s and tumor-specific OT-Is in B16-ova flank tumors. **b-c.** Gating used to identify tumor **(b)** and virus-specific **(c)** CD8s and characterize CD39 expression by residency.
